# Proposal for minimum information guidelines to report and reproduce results of particle tracking and motion analysis

**DOI:** 10.1101/155036

**Authors:** Alessandro Rigano, Caterina Strambio-De-Castillia

## Abstract

The proposed Minimum Information About Particle Tracking Experiments (MIAPTE) reporting guidelines described here aim to deliver a set of rules representing the minimal information required to report and support interpretation and assessment of data arising from intracellular multiple particle tracking (MPT) experiments. Examples of such experiments are those tracking viral particles as they move from the site of entry to the site of replication within an infected cell, or those following vesicular dynamics during secretion, endocytosis, or exocytosis. By promoting development of community standards, we hope that MIAPTE will contribute to making MPT data FAIR (Findable Accessible Interoperable and Reusable). Ultimately, the goal of MIAPTE is to promote and maximize data access, discovery, preservation, re-use, and repurposing through efficient annotation, and ultimately to enable reproducibility of particle tracking experiments. This document introduces MIAPTE v0.2, which updates the version that was posted to Fairsharing.org in October 2016. MIAPTE v0.2 is presented with the specific intent of soliciting comments from the particle tracking community with the purpose of extending and improving the model. The MIAPTE guidelines are intended for different categories of users: 1) Scientists with the desire to make new results available in a way that can be interpreted unequivocally by both humans and machines. For this class of users, MIAPTE provides data descriptors to define data entry terms and the analysis workflow in a unified manner. 2) Scientists wishing to evaluate, replicate and re-analyze results published by others. For this class of users MIAPTE provides descriptors that define the analysis procedures in a manner that facilitates its reproduction. 3) Developers who want to take advantage of the schema of MIAPTE to produce MIAPTE compatible tools. MIAPTE consists of a list of controlled vocabulary (CV) terms that describe elements and properties for the minimal description of particle tracking experiments, with a focus on viral and vesicular traffic within cells. As part of this submission we provide entity relationship (ER) diagrams that show the relationship between terms. Finally, we also provide documents containing the MIAPTE-compliant XML schema describing the data model used by Open Microscopy Environment inteGrated Analysis (OMEGA), our novel particle tracking data analysis and management tool, which is reported in a separate manuscript. MIAPTE is structured in two sub-sections: 1) Section 1 contains elements, attributes and data structures describing the results of particle tracking, namely: particles, links, trajectories and trajectory segments. 2) Section 2 contains elements that provide details about the algorithmic procedure utilized to produce and analyze trajectories as well as the results of trajectory analysis. In addition MIAPTE includes those OME-XML elements that are required to capture the acquisition parameters and the structure of images to be subjected to particle tracking.

## Introduction

A progressively larger number of systems biology “omics” studies incorporate a diverse set of data types thereby enabling multidimensional (e.g. multiple times, spaces, scales, molecules analyzed and genetic backgrounds) views of cells, organisms, or their populations. Yet it is increasingly clear that, because of the sheer complexity of biological systems, the full impact of these studies can only be realized through research involving multiple laboratories each working in parallel on the same experimental system and producing comparable results datasets. In turn, this not only requires rigorous reproducibility of results but also demands data harmonization and sharing to facilitate data re-use, re-interpretation, and meta-analysis. This level of data integration necessitates consistent generation, capture, and distribution of metadata, as well as the development of standard data structures, application programming interfaces (API), as well as tools capable of taking advantage of these shared resources.

### Minimal reporting guidelines

Scientists can correctly evaluate, interpret, and effectively re-produce image analysis results only if information describing the “origin” and “lineage” of data is recorded and made available to them. Such information is often referred to as describing “data provenance”, and it can be conceptualized as metadata capable of answering seven key questions (i.e. what, who, where, when, which, why, and how), describing events that affect transformations data undergoes during its lifetime (Ram and Liu, 2009).

With ever expanding sources of biological data, provenance metadata definition standards have become crucial to ensure that sufficient information is reported for a study to allow correct interpretation of the results, reproduction of the experiments, as well as reuse of the data. Reporting guidelines, or minimum information checklists, collectively termed Minimum Information for Biological and Biomedical Investigations (MIBBI), have been proposed as a way to understand the context, methods, data and conclusions of experiments in different domains (Taylor et al., 2008; Millard et al., 2011) and are currently maintained in the Fairsharing.org portal. Generally, these reporting guidelines are presented in three different forms: 1) Minimal reporting requirements are descriptive guidelines that make certain that sufficient metadata is provided to understand and repeat an experiment or an analysis process. 2) Controlled vocabularies ensure the use of uniform annotation to describe both experimental and algorithmic procedures, and that this annotation is at the same time human- as well as computer-readable. 3) Data formats and API standards are key to allow unhindered and straightforward access to the reported data, results and all associated metadata.

### Reporting guidelines for particle tracking

Intracellular trafficking of diffraction-limited objects such as viruses and vesicles is a fundamental process in cell biology, the understanding of which directly impacts fields as diverse as viral infection, immune regulation, metabolic disorders, and cancer. The real-time visualization of individual fluorescently-labeled intracellular vesicles and virions (i.e. bright spots) by fluorescence-microscopy, coupled with feature point tracking techniques (i.e. Single-or as recently proposed Multiple-Particle Tracking, which are abbreviated as SPT and MPT respectively; Chenouard et al., 2014) and the mathematical analysis of motion, is ideally suited to study the fate of heterogeneous populations of particles as they progress within the cell, map dynamic interactions with the underlying cellular machinery, and dissect individual transport steps (Aoyama et al., 2017; Gramlich and Klyachko, 2017; Mercer et al., 2010; Brandenburg and Zhuang, 2007; Sun et al., 2013; Li et al., 2017). As a case in point, imaging experiments following the movement of individual viral particles coupled with MPT and motion analysis have fundamentally improved understanding of the early phases of viral entry and revealed previously un-recognized entry stages (Greber and Way, 2006; Marsh and Helenius, 2006). As MPT studies become increasingly high-content and high-throughput as a result of advances in spatiotemporal microscopic resolution and algorithmic methodologies (Liberali et al., 2015; Liberali and Pelkmans, 2012; Damm and Pelkmans, 2006; Snijder et al., 2012), the need is emerging to develop: 1) solutions for both intra-and inter-lab data management; 2) infrastructure for data standardization and dissemination; and 3) approaches to facilitate the meta-analysis and interpretation of available datasets.

In order to respond to this need we set out to construct a comprehensive, open and free data exchange ecosystem for viral and vesicular trafficking data, based on the development of extensible community metadata and API standards, as well as tools capable of leveraging these standards and making them available to the bench scientist. As a first step in the direction of maximizing transparency, data harmonization, reproducibility, and re-usability of particle tracking data, we have developed a set of guidelines termed Minimum Information about Particle Tracking Experiments (MIAPTE), that follows MIBBI specifications and was deposited to Fairsharing.org in 2016 (Rigano and Strambio-De-Castillia, 2016). MIAPTE’s aim is to provide rules that detail which information should be included when reporting the results of viral and vesicular intracellular trafficking experiments while describing the MPT and motion analysis process in a fully reproducible manner. MIAPTE covers the description of the raw image data, the extraction of particle trajectories from time-series using MPT algorithms as well as the analysis of such trajectories to obtain quantitative motion descriptors from which biological understanding can be inferred. On the other hand these guidelines do not cover the generation of biological specimens and their transfer to the microscope, which might be captured by other standards (Ganzinger et al., 2012; Masuzzo, 2017; Ganzinger, 2011; Scheuermann, 2010). Thus, first and foremost, MIAPTE provides a checklist of metadata descriptors and data model specifications to describe trajectory results. In addition, MIAPTE comprises a checklist of CV terms for the minimal description of MPT analysis pipelines and results.

In order to take advantage of MIAPTE we have developed an open-access tool termed the Open Microscopy Environment inteGrated Analysis (OMEGA) for particle tracking data production, analysis, and quality control (described in a parallel manuscript; Rigano et al. in preparation), which relies on a robust, future proof repository implementing the MIAPTE schema to collect, annotate, manage and promote dissemination of these data in standardized machine and human-readable formats.

Here we present MIAPTE v0.2, the first public revision of the standard, which incorporates the experience we have accumulated during the development and testing of OMEGA as well as suggestions arising from the Cell Migration Standards Organization (CMSO) initiative (Cell Migration Standard Organization, 2014; Masuzzo and Martens, 2015). MIAPTE v0.2 is intended to be a Request for Comments (Crocker and Postel, 2008). Members of the scientific and developer communities are invited to collaborate with us to modify, extend, and improve the model.

## Results

MIAPTE aims at standardizing the process used to extract particle trajectory data from time series and to analyze their motion as well as providing community guidelines for reporting trajectory data and motion analysis results. In order to serve the needs of a diverse set of possible users, MIAPTE guidelines are provided in three formats: 1) a glossary in tabular format, where data elements are presented together with their properties, their definitions, their cardinality, requirement level (http://www.ietf.org/rfc/rfc2119.txt), as well as recommended sources for their annotation, examples, and notes (Supplementary information 1, Table I: Glossary of MIAPTE guidelines). 2) ER diagrams, which depict each entity together with its attributes as well as the relationship between one another (Figures 2 and 3). 3) XML schemas, which were created based on our implementation of the MIAPTE guidelines in OMEGA. Such schema can be used to generate the corresponding data structures (Supplementary information 2 and 3, MIAPTE xsd schema files).

**Figure 1:**
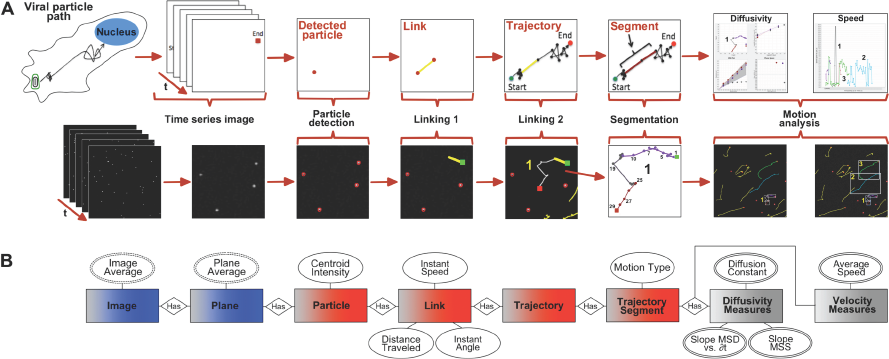
A typical workflow for tracking multiple intracellular particles and analyzing their motion. A) Schematic representation a typical workflow for the estimation of the sub-cellular trajectories taken by individual particles and for the computation of biologically meaningful measures from their coordinates While, the example in the figure represents a viral tracking experiment, the same workflow can be used for any other diffraction limited intracellular object. After **<u>Particle Detection</u>** identifies individual fluorescent local-maxima and estimates their sub-pixel centroid coordinates in each time frame, **<u>Particle Linking</u>** generates trajectories by linking the position of each bright spot in one time frame with its position in subsequent frames. In order to interpret the biological significance particle movement, first **<u>Trajectory segmentation</u>** is used to subdivide trajectories into uniform mobility; then **<u>Motion analysis</u>** is used to extract mobility and diffusivity measures capable of describing the spatiotemporal dynamic behavior of individual viral particles. B) Entity Relationship diagram representing the corresponding OME-XML (Blue) MIAPTE (Red and Grey) elements utilized to capture data pertaining to each step of the data-flow. Multi-values, derived attributes (double-lined, dashed ovals) are represented here as examples of aggregate measures that could optionally be associated at the Image, Dataset or Project levels.

**Figure 2:**
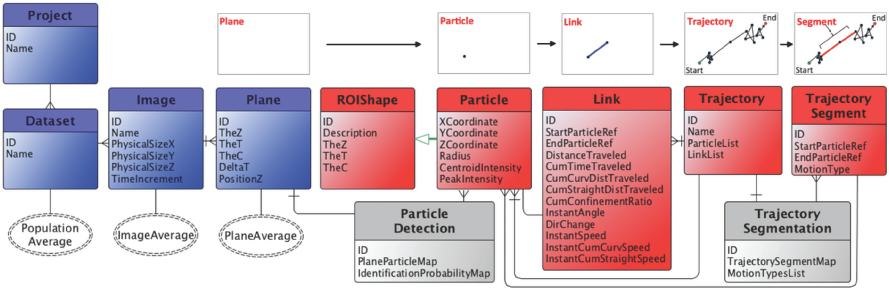
Section 1 of Minimum Information About single Particle Tracking Experiments (MIAPTE) standard proposal. Entity-Relationship diagram representing the Trajectory Elements section of MIAPTE. Red entities represent MIAPTE specific elements and attributes responsible for storing information related to single particle tracking data. Blue entities represent elements shared with the OME-XML model (http://www.openmicroscopy.org/Schemas/Documentation/Generated/OME-2016-06/ome.html) that are directly related to MIAPTE. Grey entities represent elements shared with the Analysis Elements section of MIAPTE. Multi-values, derived attributes (double-lined, dashed ovals) are represented here as examples of aggregate measures that could optionally be associated at the Image, Dataset or Project levels. OME-XML: ExperimenterGroup, OME-XML: Experimenter and unit attributes have been omitted for simplicity sake. See main text and Supplementary Table I for more explanations on individual elements and attributes.

**Figure 3:**
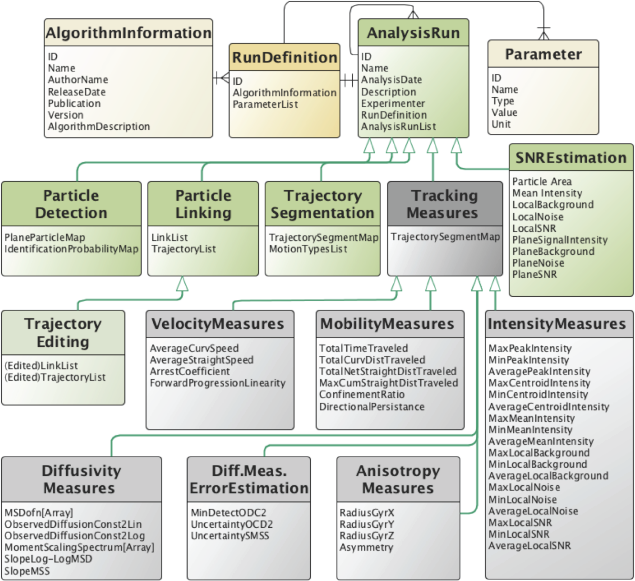
Section 2 of Minimum Information About single Particle Tracking Experiments (MIAPTE) guidelines proposal. Entity-Relationship diagram representing Section 2 of MIAPTE, which contains elements responsible for storing analysis definition metadata and analysis results. Yellow elements represent classes responsible for storing information relative algorithms used for each analysis step. Green elements represent elements that extend the Analysis Run element. Grey elements are Tracking Measures elements and their extensions. See main text and Supplementary Table I for more explanations on individual elements and attributes.

In order to avoid duplication, MIAPTE encapsulates OME-XML, which is considered the current standard for capturing and sharing metadata describing microscopic acquisition parameters as well as the characteristics of multidimensional image data (Figure 2, blue elements; (Goldberg et al., 2005; Linkert et al., 2010a)). At the same time MIAPTE extends OME-XML to capture particle tracking and motion analysis processes and results downstream of raw-images. Specifically, the MIAPTE guidelines are subdivided in two sub-sections (Figures 2 and 3):

1. Section 1 contains Trajectory Elements (Figure 2, red elements), which capture information about the output of MPT.
2. Section 2 contains Motion Analysis Elements (Figure 3), which capture information about the tracking and motion analysis workflow utilized to produce trajectories (grey elements) and motion analysis results associated with such trajectories (yellow and green elements).

All MIAPTE elements store a Unique Identifier for every data element. In addition, MIAPTE includes OME-XML ExperimenterGroup and extends Experimenter so that it can be put in direct “has-a” relationship to specific analysis runs and results.

### Section 1: Trajectory Elements

The Trajectory Elements section of MIAPTE contains semantic data types that are required to fully describe the minimal set of results arising from MPT and motion analysis workflows (Figure 2, red and grey elements).

This section includes the following entities and their attributes: Particle, Link, Trajectory and Segment. Particle extends ROIShape, an element introduced to capture a fusion of the OME-XML ROI and Shape element (Goldberg et al., 2005; Linkert et al., 2010a), and represents bright spots identified by the particle detection algorithm. Link models a connection between two particles while Trajectory contains a list of particles and their links. Segment represents a portion of a Trajectory, which is characterized by a uniform motion type. To model this concept, each TrajectorySegment has an associated MotionType.

### Section 2: Analysis Elements

The Analysis Elements section of the model (Figure 3) contains components that describe the motion analysis results and the process that generated them. Captured information includes algorithm description, parameters definition as well as related algorithm-specific or parameter-dependent analysis results. For example, in our OMEGA implementation of this model, algorithms obligatorily provide the set of information that is defined in the AlgorithmInformation entity. This is then used to describe RunDefinition, which is linked to a specific AnalysisRun and to a series of entities defining the state of associated Parameters. The generic entity AnalysisRun includes a timestamp, is linked to an Experimenter, and might contain a name, if given. It might also be linked to a series of dependent AnalysisRun elements if that is required to correctly capture different workflow steps and their corresponding results. AnalysisRun is extended by a series of elements, which are responsible to capture the output of each set of executed runs.

## Summary

Defining this common set of metadata to guide researchers in reporting scientific context is important to facilitate results reproducibility and re-use. This in turn can lead to more robust data repositories, more reliable journal publications, and a better way to assess the quality of scientific output, which is in turn essential for funding purposes.

The MIAPTE minimum reporting guidelines (Rigano and Strambio-De-Castillia, 2016) specify the degree of detail that needs to be captured, for reporting MPT analysis workflows as well as the results arising from them. Importantly, MIAPTE could be considered an extension of the OME-XML metadata definition standard (Goldberg et al., 2005; Linkert et al., 2010b), in as far as elements, attributes, and types defined in that model are included here when appropriate to avoid "reinventing the wheel". In addition by conforming to minimum information criteria currently maintained by Fairsharing.org (Taylor et al., 2008), the use of MIAPTE will enable both the effective interpretation and assessment of MPT data, and, importantly, the reproduction of the analysis process that generated it.

As for all MIBBI standards, MIAPTE’s guidelines are expected to evolve. In order to start the process of refinement and extension of the guidelines we ask that all relevant members of the MPT community (i.e. experimental biologists, algorithm developers, software engineers and data scientist) test the proposed metadata checklist and provide feedback for their use and improvement. We hope that this process will allow us to reach agreement on the minimal descriptors required to understand, interpret, evaluate, replicate, and disseminate cell migration experiments. To contribute, please browse to the website at: http://big.umassmed.edu/omegaweb/resources/miapte/. Alternatively you can comment on the FairSharing.org website at: https://fairsharing.org/feedback/standard/2066.

## Conclusions

We hope that this MPT checklist will serve as a common denominator to guide experimental design, capture important parameters, and be used as a standard format for stand-alone data publications. In turn the ability of publishing datasets will create efficient linkages between data and knowledge extraction, importantly allowing for appropriate attribution to data generators and infrastructure science builders in the post-genomics era.

## Supplemental Material

**Supplementary information 1 -- Table I: Glossary of MIAPTE guidelines**

https://github.com/QmegaProiect/MIAPTE/blob/master/2017-07-12v02/2017-07-12_MIAPTE_Guidelines_v02.xlsx

**Supplementary information 2 -- MIAPTE Section 1 xsd schema**

https://github.com/OmegaProject/MIAPTE/blob/master/2017-07-12_v02/2017-07-12MIAPTESection1rajectoryElementsv02.xsd

**Supplementary information 3 -- MIAPTE Section 2 xsd schema**

https://github.com/OmegaProject/MIAPTE/blob/master/2017-07-12_v02/2017-07-12_MIAPTE_Section_2_AnalysisElements_v02.xsd

